# ADP-heptose is a pathogen-associated molecular pattern of *Shigella flexneri* infection

**DOI:** 10.1101/407262

**Authors:** Diego García-Weber, Anne-Sophie Dangeard, Johan Cornil, Linda Thai, Héloise Rytter, Laurence A. Mulard, Cécile Arrieumerlou

## Abstract

During an infection, the detection of pathogens is mediated through interactions between pathogen-associated molecular patterns (PAMPs) and pathogen recognition receptors. β-Heptose 1,7-bisphosphate (βHBP), a metabolite of the lipopolysaccharide (LPS) biosynthesis pathway, was recently identified as a PAMP of gram-negative bacteria. It was reported that βHBP sensing leads within minutes to oligomerization of the protein TIFA, a mechanism controlling NF-κB activation and pro-inflammatory gene expression. Here, we compared the ability of chemically synthesized βHBP and *Shigella flexneri* lysate to induce TIFA oligomerization in epithelial cells. In contrast to lysate, we found that βHBP fails to trigger rapid oligomerization of TIFA. βHBP only induces delayed signaling, suggesting that it has to be processed intracellularly to induce inflammation. By dissecting the LPS biosynthesis pathway with deletion mutants and functional complementation experiments, we show that ADP-D-*glycero*-β-D-*manno*-heptose and ADP-L-*glycero*-β-D-*manno*-heptose are the bacterial metabolites responsible for rapid TIFA oligomerization, and that they strongly induce interleukin-8 expression during *S. flexneri* infection. Altogether, our results rule out a major role of βHBP in *S. flexneri* infection and identify ADP-heptose as a new PAMP.

## INTRODUCTION

During an infection, an immune response is triggered by the detection of pathogens. This is mainly achieved by cells of the innate immune system including macrophages and neutrophils. Epithelial and endothelial cells are also involved in this process. They act as sentinels of the immune system and contribute, via the secretion of inflammatory mediators, to elicit an integrated immune response. At the molecular level, pathogen recognition is mediated by the interactions between pathogen-associated molecular patterns (PAMPs) and pathogen recognition receptors. Well characterized bacterial PAMPs include among others lipopolysaccharide (LPS), flagellin, peptidoglycan, lipoteichoic acid and DNA. β-Heptose 1,7-bisphosphate (βHBP) was recently identified as a PAMP of *Neisseria meningitidis* [1]. βHBP is a soluble metabolite of the LPS biosynthesis pathway. In *N. meningitidis*, it is produced by the D-β-D-heptose 7-phosphate kinase activity of the HldA protein that phosphorylates D-*glycero*-β-D-*manno*-heptose 7-phosphate into βHBP [2]. Gaudet et al. showed that it can be secreted by *N. meningitidis* and sensed in the cytosol of eukaryotic cells after being internalized by endocytosis [1]. Our laboratory reported that βHBP sensing was involved in the inflammatory response of epithelial cells to *Shigella flexneri* and *Salmonella typhimurium* infection [3]. This finding was based on the analysis of the *hlde* and *gmhb* deletion mutants of the LPS biosynthesis pathway. In these bacterial species closely related to *Escherichia coli, hlde* encodes a bifunctional protein that harbors both D-β-D-heptose 7-phosphate kinase and D-β-D-heptose 1-phosphate adenylyltransferase activities, responsible for the production of βHBP and ADP-D-*glycero*-β-D-*manno*-heptose (ADP-D-β-D-heptose), respectively [4]. The dephosphorylation of βHBP into D-*glycero*-β-D-*manno* heptose 1-phosphate is provided by the phosphatase activity of GmhB [5]. We showed that infection with an *hlde* deletion (Δ*hldE*) mutant failed to induce the expression of the inflammatory cytokine interleukin-8 (IL-8) whereas this latter was restored after infection with a mutant deleted for *gmhb* (Δ*gmhB)* [3]. *S. flexneri* infection leads to oligomerization of the proteins TRAF-interacting protein with FHA domain-containing protein A (TIFA) and TNF receptor-associated factor 6 (TRAF6), and activation of the transcription factor NF-κB. TIFA is a 20-kDa protein that was first identified as a TRAF6-interacting protein in a yeast two-hybrid screen [6]. It contains a forkhead-associated (FHA) domain, known to bind phosphothreonines and phosphoserines [7], and a consensus TRAF6-binding motif. TIFA oligomerization is dependent on the phosphorylation of a threonine at position 9 and on the FHA domain [8]. Unphosphorylated TIFA is thought to exist as an intrinsic dimer constitutively linked to TRAF6. When T9 is phosphorylated, this is recognized by the FHA domain of other TIFA dimers leading to its oligomerization and thus the oligomerization of TRAF6. This enhances the E3 ubiquitin ligase activity of TRAF6 and leads to the activation of the NF-κB pathway [9]. We recently revealed the role of ALPK1 in bacterial infections and showed that this atypical kinase of the α-kinase family regulates the oligomerization of TIFA [3]. More recently, the role of βHBP sensing and the contribution of the ALPK1/TIFA pathway were also observed in *Helicobacter pylori* infection [10–12], reinforcing the broad implication of this newly emerging pathway in innate immunity. In addition to the use of LPS biosynthesis mutants, the pro-inflammatory activity of βHBP was directly confirmed. It was first synthesized *in vitro* from the sedoheptulose 7-phosphate precursor by the sequential activities of purified GmhA and HldE [1]. More recently, βHBP was also chemically synthesized [13–16]. Its pro-inflammatory activity was validated on the basis that it induced NF-κB activation after several hours of treatment [14,15]. In this report, we compared the ability of chemically synthesized βHBP and *S. flexneri* lysates to induce the oligomerization of TIFA and the production of IL-8 in human epithelial cells. In contrast to lysate from wild-type bacteria, we found that βHBP was unable to induce rapid oligomerization of TIFA. Oligomerization was only observed after more than one hour, indicating that cellular processing of βHBP was necessary to trigger this mechanism. It also suggested that another PAMP was therefore responsible for the early mechanism of TIFA oligomerization occurring during infection [3]. By combining deletion mutants and functional complementation experiments, we identified ADP-heptose as a new PAMP of *S. flexneri* infection.

## RESULTS AND DISCUSSION

### βHBP fails to induce rapid oligomerization of TIFA

In order to characterize the role of βHBP in innate immunity, this bacterial metabolite was chemically synthesized. A synthetic route was implemented, that was inspired, for the most part, from the work of Vincent and coworkers [17] (Figure 1 and Materials and Methods). Briefly, starting from thiophenyl α-D-mannoside (**2**) [18], easily synthesized in three steps from D-mannose (**1**), a sequence of protection/deprotection steps afforded the production of 2,3,4-tri-*O*-benzylated derivative **3** on a large scale (55 mmol) and in good yield. Homologation of this thiomannoside intermediate by means of a Parikh-Doering oxidation/Wittig sequence gave the terminal olefin-containing heptoside **4** [19]. The latter was subsequently reacted with catalytic osmium tetroxide to obtain a 7:3 diastereomeric mixture of diols, from which the major isomer was isolated by chromatography. A subsequent four step protection/deprotection procedure was applied to give hemiacetal **5**. The hemiacetal was submitted to a Mitsunobu-mediated *bis*-phosphorylation at OH-1 and OH-7 by use of a combination of dibenzyl phosphate, DIAD, P(ClPh)3, and triethylamine in tetrahydrofuran [13]. Chromatography of the obtained α/β mixture of the perbenzylated diphosphorylated monosaccharide gave the α-anomer and β-anomer in 30% and 41% yield, respectively. The purified β-heptose was subjected to a palladium-catalyzed hydrogenation followed by H^+^/Na^+^ exchange to give the expected βHBP [13]. The pro-inflammatory activity of chemically synthesized βHBP was first tested by measuring the production of IL-8 after transient permeabilization of cell membranes with digitonin. In this assay, the cellular process of βHBP internalization is bypassed and the mechanisms of PAMP recognition and inflammatory signaling can thus be directly assessed. HeLa cells were treated for 30 minutes with digitonin and synthetic βHBP at indicated concentrations (Figure 2A and 2B). As a control, cells were treated with digitonin alone or digitonin and wild-type (wt) *S. flexneri* bacterial lysate. Cells were then washed, treated with monensin to trap IL-8 intracellularly, and IL-8 was monitored in single cells after 6 hours with automated microscopy (see Materials and Methods). Data showed that synthetic βHBP induced the production of IL-8 in a dose-dependent manner (Figure 2A and 2B). Interestingly, the βHBP concentration required to induce IL-8 expression to the level of *S. flexneri* lysate was unexpectedly high (Figure 2A and 2B). Since a millimolar-range concentration of βHBP is probably not reached in our bacterial lysate [20], this observation raised the possibility that additional bacterial factors may contribute to elicit the IL-8 response. βHBP-induced IL-8 expression was previously shown to depend on the oligomerization of TIFA proteins, a process controlling NF-κB activation [1,3]. We therefore analyzed the effect of chemically synthesized βHBP on TIFA oligomerization. HeLa cells expressing GFP-tagged TIFA protein (TIFA-GFP) were treated as described above and TIFA-GFP oligomerization was monitored after 6 hours. It was quantified in single cells by automated image analysis as described in Materials and Methods. In line with IL-8 data, βHBP induced the formation of large TIFA-GFP oligomers in a dose-dependent manner (Figure 2C and 2D). This result was in agreement with previous reports showing that synthetic βHBP induces NF-κB activation after several hours of treatment [14,15]. Given that TIFA oligomerization is observed within minutes of *S. flexneri, S. typhimurium* or *H. pylori* infections [3,10], we also monitored TIFA oligomerization after 30 minutes of treatment. Strikingly, whereas cells treated with *S. flexneri* lysate induced massive oligomerization of TIFA-GFP within this time period, synthetic βHBP failed to do so (Figure 2E). At least one hour was required to observe significant oligomerization of

**Figure 1:**
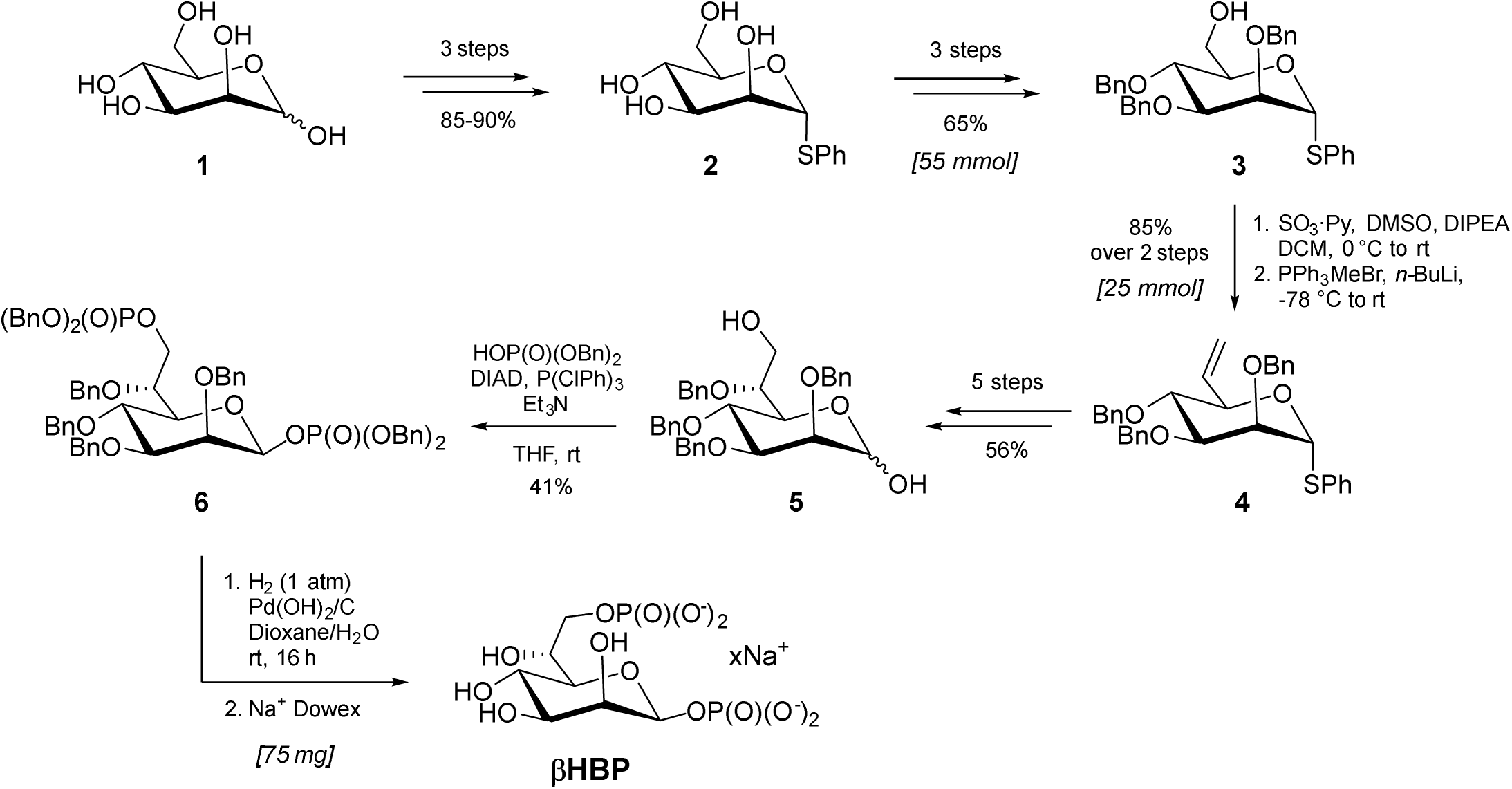
Chemical synthesis of βHBP. Chemical synthesis of βHBP from D–mannose (1): overview.

**Figure 2:**
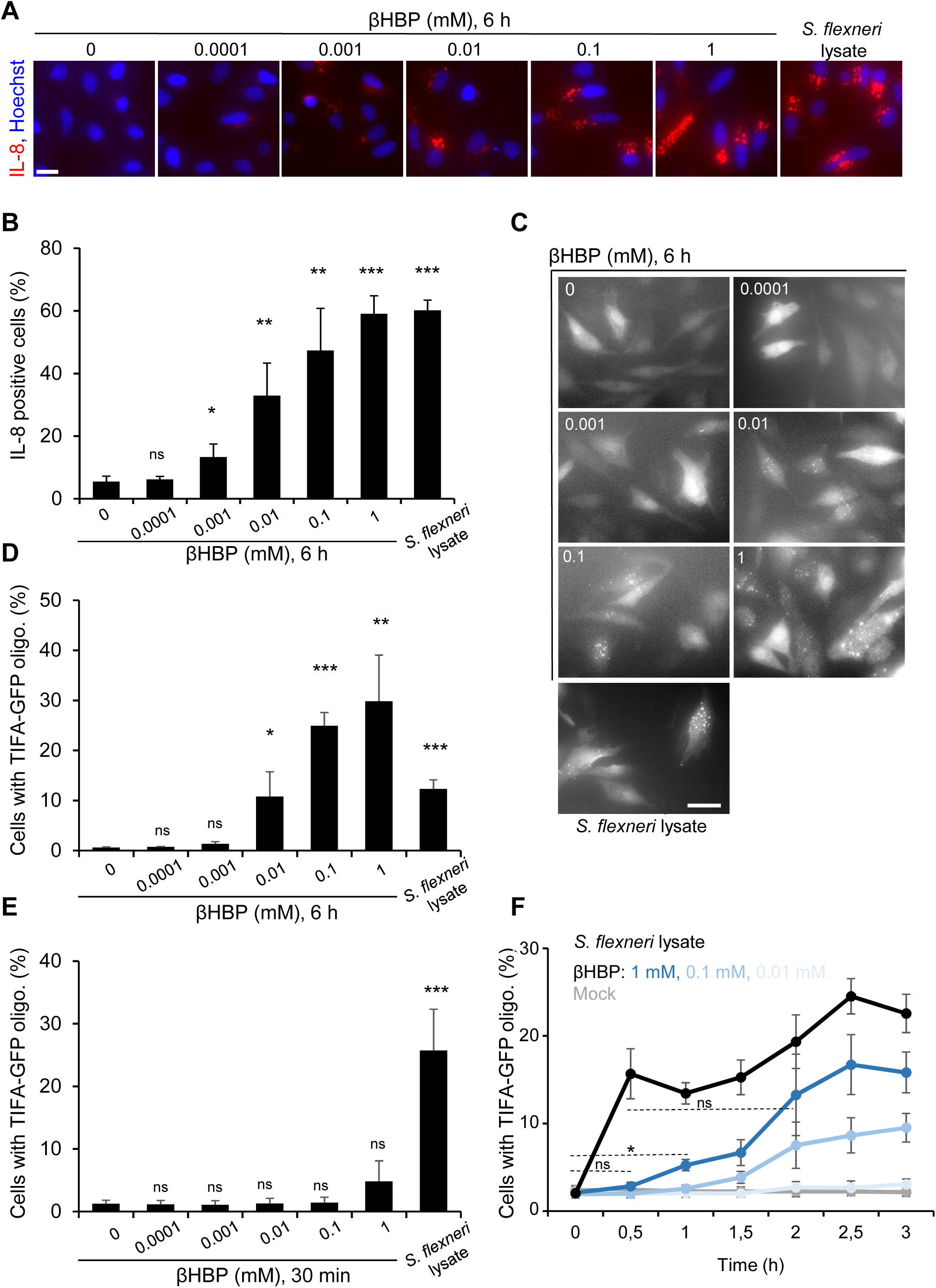
βHBP fails to induce rapid oligomerization of TIFA. **A**) βHBP-induced IL-8 production is dose-dependent. HeLa cells were treated for 30 min with digitonin alone, digitonin and wild-type *S. flexneri* lysate or digitonin and βHBP at indicated concentrations. IL-8 production was monitored by immunofluorescence microscopy after 6 h. Hoechst and IL-8 are shown in blue and red, respectively. Scale bar: 20 μm. **B**) Quantification of IL-8 measurements. Cells were treated as in A and IL-8 was quantified by automated image analysis. Data represent the mean +/-SD of 3 independent experiments. **C**) TIFA oligomerization is visible after 6 h of βHBP treatment. HeLa cells expressing TIFA-GFP were stimulated as in A, fixed and observed by epifluorescence microscopy 6 h post treatment. Scale bar, 20 μm. **D**) Quantification of TIFA oligomerization as shown in C. Cells with TIFA-GFP oligomers were quantified by automated image analysis. Data correspond to the mean +/-SD of 3 independent experiments. **E**) βHBP fails to induce rapid oligomerization of TIFA-GFP. HeLa cells expressing TIFA-GFP were stimulated as in A, fixed and observed by microscopy 30 min post treatment. Cells with TIFA-GFP oligomers were quantified by automated image analysis. Data correspond to the mean +/-SD of 3 independent experiments. **F**) Kinetics of TIFA-GFP oligomer formation upon βHBP treatment. Cells were stimulated as in A and TIFA-GFP oligomerization was monitored at indicated time points. Cells with TIFA-GFP oligomers were quantified by automated image analysis. Data correspond to the mean +/-SEM of 4 independent experiments. For comparison between mock and treated conditions (B, D, E, F) or *S. flexneri* lysate and βHBP (F, dashed bar), statistical significance was assessed using unpaired, two-tailed Student’s *t*-test, p*<0.05, p**<0.005, p***<0.0005, non-significant (ns).

TIFA-GFP in a small fraction of cells treated with 1 mM synthetic βHBP (Figure 2F). For this latter concentration, the level of TIFA-GFP oligomerization observed in cells treated with *S. flexneri* lysate was only reached after 2 hours of βHBP treatment (Figure 2F). Such delay in the response to βHBP suggested that this bacterial metabolite has to be processed intracellularly to trigger inflammation signaling, and that the sensing of another bacterial PAMP is likely responsible for the rapid mechanism of TIFA oligomerization observed upon *S. flexneri* lysate treatment.

### Bacterial ADP-heptose synthesis is required for rapid oligomerization of TIFA

Our results showing that βHBP was unable to trigger rapid oligomerization of TIFA-GFP (Figure 2E and 2F) opened a new avenue for the identification of new bacterial PAMPs inducing inflammatory signaling upon *S. flexneri* infection. Given that we previously reported that Δ*hldE S. flexneri* bacteria completely failed to trigger TIFA oligomerization and inflammatory gene expression [3], we further analyzed the role of the bifunctional enzyme HldE. More specifically, we analyzed the contribution of its D-β-D-heptose 7-phosphate kinase and D-β-D-heptose 1-phosphate adenylyltransferase activities that synthesize βHBP and ADP-D-β-D-heptose, respectively (Figure 3A). For this, we took advantage of the fact that in *N. meningitidis*, these enzymatic activities are harbored by two distinct proteins encoded by the *hlda* and *hldc* genes [2], and examined the ability of lysates from Δ*hldE S. flexneri* complemented with *hlde, hlda, hldc* or *hlda* and *hldc* to induce TIFA oligomerization in digitonin-permeabilized cells. Whereas treatment with wt *S. flexneri* lysate induced massive oligomerization of TIFA-GFP at 30 minutes, lysate from Δ*hldE* bacteria failed to do so (Figure 3B). As expected, complementation of Δ*hldE* bacteria with a plasmid encoding HldE (pHldE) fully restored fast oligomerization of TIFA, confirming the critical role of HldE in this process. Interestingly, lysate from Δ*hldE* bacteria complemented with plasmid-encoded HldA (pHldA) had no effect at 30 minutes (Figure 3B). This result was in line with synthetic βHBP data showing that this metabolite was indeed unable to trigger rapid oligomerization of TIFA-GFP (Figure 2E and 2F). By showing that bacterial D-β-D-heptose 1-phosphate adenylyltransferase activity was required, this new finding indicated that the PAMP eliciting rapid oligomerization was produced downstream of the adenylyltransferase enzymatic reaction within the LPS biosynthesis pathway (Figure 3A). Complementation of Δ*hldE S. flexneri* with plasmid-encoded HldC (pHldC) failed also to trigger TIFA oligomerization at 30 minutes, showing that bacterial synthesis of βHBP, although not sufficient, was required to trigger early signaling. As expected, this process was restored with Δ*hldE* bacteria complemented with both pHldA and pHldC (Figure 3B). The observation that double complementation only led to partial restoration may be explained by a low expression of the HldA and HldC proteins encoded from two different plasmids, or by a difference in enzymatic efficacy between HldE of one part and HldA and HldC on the other hand. Together, these results showed that D-β-D-heptose 7-phosphate kinase and D-β-D-heptose 1-phosphate adenylyltransferase activities are both required to stimulate rapid pro-inflammatory signaling, indicating that bacterial synthesis of βHBP and ADP-D-β-D-heptose is necessary. In order to further characterize the PAMP involved in rapid inflammatory signaling, we analyzed the *hldD* and *waaC* deletion mutants of the LPS biosynthesis pathway, named Δ*hldD* and Δ*waaC* respectively. HldD harbors an epimerase activity catalyzing the interconversion between ADP-D-β-D-heptose and ADP-L-*glycero*-β-D-*manno*-heptose (ADP-L-β-D-heptose) [21] (Figure 3A). Directly downstream of HldD, WaaC, which has an ADP-heptose-lipopolysaccharide heptosyltransferase activity, allows heptose transfer to the LPS core [21]. Data showed that Δ*hldD* and Δ*waaC* lysates were both very effective at triggering TIFA-GFP oligomerization at 30 minutes, indicating that the PAMP controlling this process was produced upstream of HldD or WaaC respectively, and was accumulated in these mutants. Altogether, by showing that the PAMP of interest was dependent on βHBP, produced downstream of the D-β-D-heptose 1-phosphate adenylyltransferase enzymatic reaction, and upstream of HldD or WaaC (Figure 3B), our results point to the role of ADP-D-β-D-heptose and ADP-L-β-D-heptose (ADP-D/L-β-D-heptose, also named ADP-heptose for simplification reason) in triggering rapid oligomerization of TIFA upon treatment with *S. flexneri* lysate. TIFA-GFP oligomerization and IL-8 production were also analyzed at 6 hours. Based on synthetic βHBP data (Figure 2A, 2B, 2C, 2D and 2F), we hypothesized that at this time point βHBP and ADP-D/L-β-D-heptose contribute to inflammatory signaling. Our data confirmed a critical role for bacterial expression of HldE in lysate-induced TIFA-GFP oligomerization and IL-8 production at 6 hours (Figure 3C and 3D). In line with synthetic βHBP data (Figure 2A, 2B, 2C and 2D), pHldA-mediated complementation partially restored TIFA-GFP oligomerization and IL-8 production (Figure 3C and 3D), showing that bacterial synthesis of βHBP is sufficient to trigger delayed inflammatory responses. Complementation of Δ*hldE S. flexneri* lysate with pHldC failed to trigger TIFA-GFP oligomerization (Figure 3C) as well as IL-8 production (Figure 3D), showing that bacterial synthesis of βHBP is strictly required for both processes. Complementation with pHldA and pHldC partially restored both oligomerization and IL-8 expression (Figure 3C and 3D). The observation that expressing HldA or both HldA and HldC in Δ*hldE* bacteria had a rather similar effect may be explained by a higher expression of HldA in single complementation experiments and enhanced βHBP concentration in bacteria lacking D-β-D-heptose 1-phosphate adenylyltransferase activity. As expected, lysates from Δ*hldD* and Δ*waaC* mutants induced massive TIFA-GFP oligomerization and IL-8 production, confirming the role of ADP-D/L-β-D-heptose in inflammation (Figure 3C and 3D). Altogether, our data showed that βHBP is only able to trigger delayed inflammatory signaling, suggesting that it is not directly sensed by the innate immune system but that it first has to be processed in cells. In addition, our results reveal that the bacterial factors ADP-D/L-β-D-heptose trigger early oligomerization of TIFA-GFP and significantly contribute to the inflammatory response observed upon treatment with *S. flexneri* lysate.

**Figure 3:**
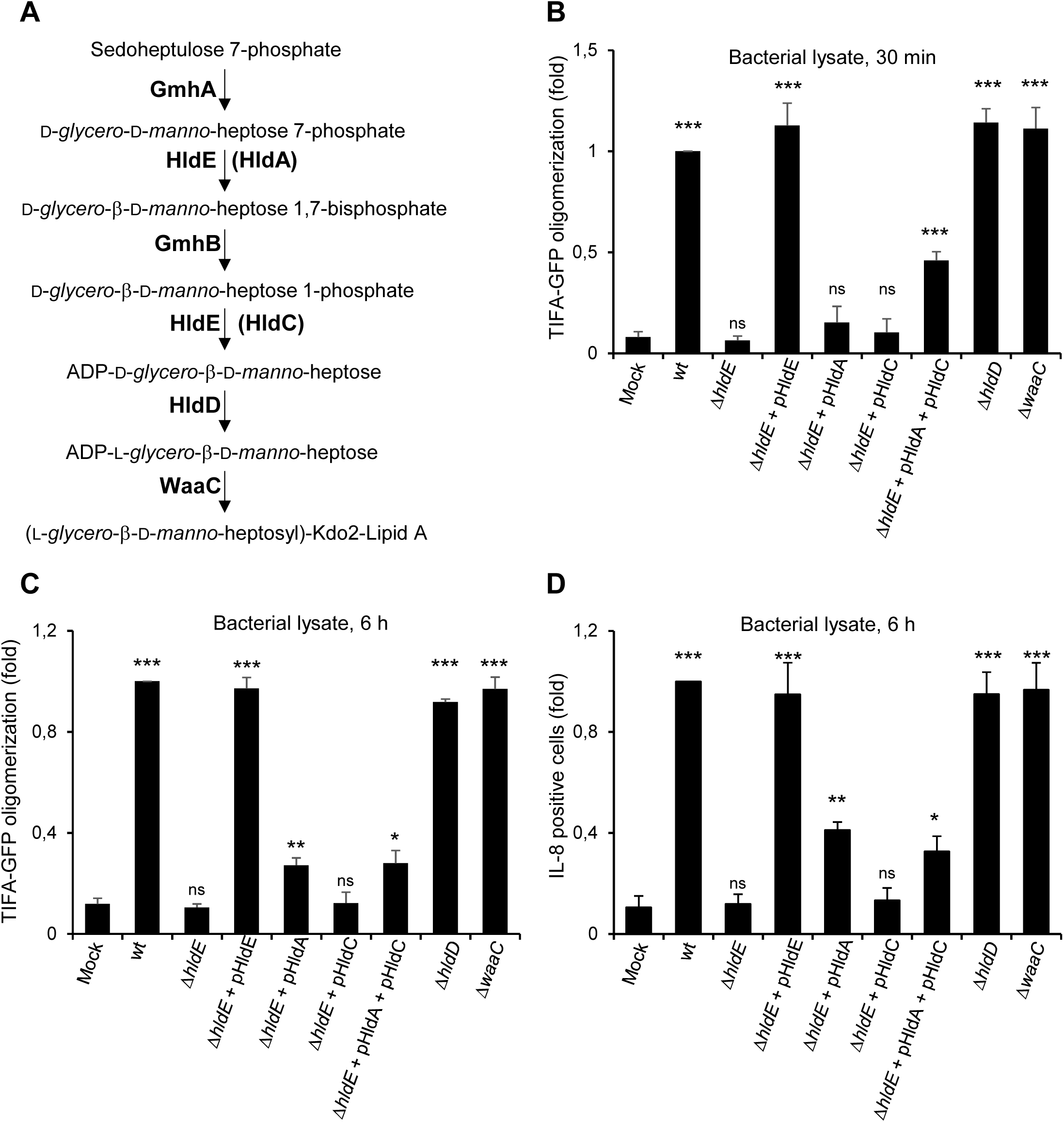
Bacterial ADP-heptose synthesis is required for rapid oligomerization of TIFA A) ADP-heptose biosynthetic pathway. HldA and HldC from *N. meningitidis* are shown in brackets. **B**) Quantification of TIFA-GFP oligomerization at 30 min. HeLa cells were treated with digitonin alone (Mock) or digitonin and a lysate from indicated *S. flexneri* strains for 30 min. Cells with TIFA-GFP oligomers were quantified by automated image analysis. Data were normalized to wt lysate treatment. Data correspond to the mean +/-SD of 3 independent experiments. **C**) Quantification of TIFA-GFP oligomerization at 6 h. HeLa cells were treated with digitonin alone (Mock) or digitonin and a lysate from indicated *S. flexneri* strains for 30 min and washed. The fraction of cells with TIFA-GFP oligomers was quantified 6 h post treatment. Data were normalized to wt lysate treatment. Data correspond to the mean +/-SD of 3 independent experiments. **D**) Quantification of IL-8 production at 6 h. HeLa cells were treated with digitonin alone (Mock) or digitonin and a lysate from indicated *S. flexneri* strains for 30 min and washed. The fraction of IL-8-positive cells was quantified 6 h post treatment. Data were normalized to wt lysate treatment. Data correspond to the mean +/-SD of 3 independent experiments. For comparison between mock and treated conditions (B, C, D), statistical significance was assessed using unpaired, two-tailed Student’s *t*-test, p*<0.05, p**<0.005, p***<0.0005, non-significant (ns).

### ADP-heptose is a PAMP of *S. flexneri* infection

The use of digitonin-permeabilized cells is a powerful method to directly address PAMP recognition and inflammatory signaling. However, it does not provide information on the actual involvement of a given PAMP during infection. The respective contribution of βHBP and ADP-D/L-β-D-heptose was therefore tested in *S. flexneri* infection experiments. For this purpose, HeLa cells were infected with wt *S. flexneri* or with the different mutants of the LPS biosynthesis pathway described above, and TIFA-GFP oligomerization was visualized 30 minutes post infection. Representative images are shown in Figure 4A. As previously reported [3], wt bacteria induced rapid oligomerization of TIFA-GFP in infected cells. In contrast, the Δ*hldE* mutant failed to do so whereas pHldE-complementation fully restored oligomerization. Together, these results confirmed the critical implication of HldE in the early host response to *S. flexneri* infection. In line with data obtained with bacterial lysates (Figure 3B), complementing the Δ*hldE* mutant with pHldA or pHldC failed to trigger TIFA-GFP oligomerization at 30 minutes (Figure 4A), showing that bacterial D-β-D-heptose 7-phosphate kinase and D-β-D-heptose 1-phosphate adenylyltransferase activities are both necessary to trigger rapid oligomerization of TIFA-GFP. This mechanism is indeed restored when cells were infected with pHldA and pHldC-complemented Δ*hldE* bacteria (Figure 4A). Finally, oligomerization was observed in cells infected with Δ*hldD* and Δ*waaC* mutants, showing that the PAMP responsible for this process was produced upstream of HldD or WaaC, respectively. In *E. coli*, Δ*hldE* and Δ*waaC* mutants have a phenotype of increased bacterial auto-aggregation and stronger cell surface hydrophobicity compared to wt bacteria [22]. In *S. flexneri*, we previously observed that these mutants are at least hundred times more invasive than wt [3], making the comparison of TIFA oligomerization between all strains inappropriate. For this reason, only mutants that had the same range of infectivity (Figure 4B) were considered for quantification. Data confirmed that none of the single-complemented mutants induced oligomerization in infected cells (Figure 4C). In contrast, although infectivity of the double-complemented mutant was slightly reduced compared to Δ*hldE* bacteria (Figure 4B), it significantly led to TIFA-GFP oligomerization in infected cells (Figure 4C). The weak impact of HldA and HldC expression on bacterial infectivity compared to the hundred fold change that was previously observed between Δ*hldE* and wt [3] suggested that the LPS of these bacteria was still significantly altered and that complementation was therefore only partial. This observation was correlated with the rather modest restoration of inflammatory signaling observed with the double-complemented mutant (Figure 3B, 3C and 3D), and may be explained by low expression or weak enzymatic activities of HldA and HldC in these bacteria. In line with lysate data (Figure 3B), infection with Δ*hldD* and Δ*waaC* mutants induced massive oligomerization at 30 minutes (Figure 4C). Altogether, these results showed that βHBP is not sufficient to trigger early signaling in *S. flexneri* infected cells and revealed the critical role of ADP-D/L-β-D-heptose in this process. In order to better understand the dynamics of the inflammatory response during infection, TIFA-GFP oligomerization and IL-8 production were also analyzed 4 hours post infection. In agreement with a role of βHBP in late signaling, oligomerization was visible in a small fraction of cells infected with pHldA-complemented bacteria (Figure 4D). However, no IL-8 was observed in response to this infection (Figure 4E). We hypothesized that the level of TIFA-GFP oligomerization was too low to elicit IL-8 production within 4 hours. This time point was chosen to prevent cell toxicity due to bacterial replication. As expected, infection with pHldC-complemented bacteria failed to trigger TIFA-GFP oligomerization (Figure 4D) and IL-8 production (Figure 4E), confirming the strict requirement of βHBP bacterial synthesis. Infection with the double-complemented mutant induced significant oligomerization of TIFA and IL-8 production, showing that the presence of both enzymes allows IL-8 expression. This latter is more likely induced by ADP-D/L-β-D-heptose than βHBP since, within 4 hours of infection, the single pHldA-complemented mutant was unable to produce IL-8 (Figure 4E). Finally, infection with Δ*hldD* and ΔwaaC mutants showed massive oligomerization and IL-8 production, confirming the ability of ADP-D/L-β-D-heptose to trigger inflammation during *S. flexneri* infection.

**Figure 4:**
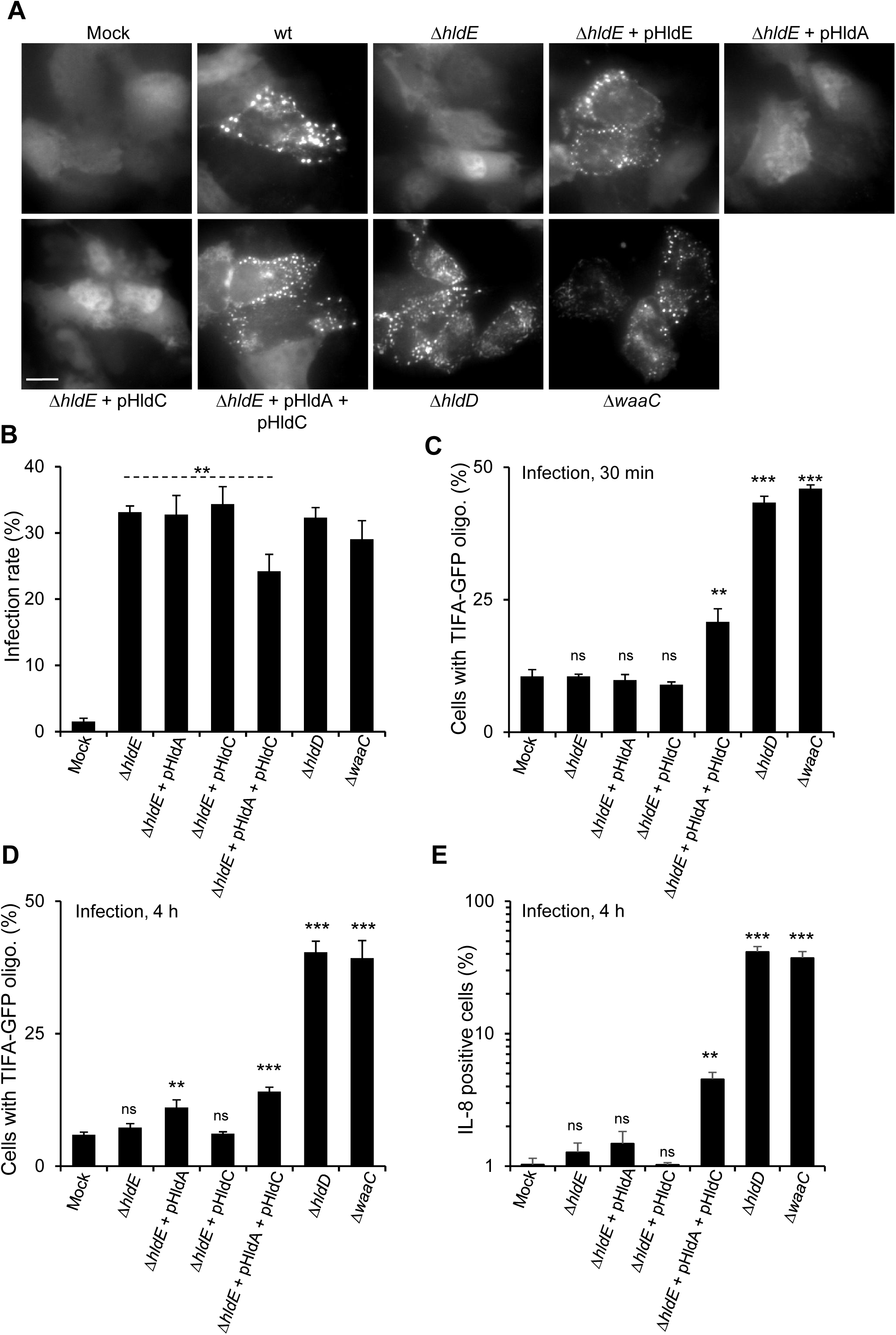
ADP-heptose is a PAMP of *S. flexneri* infection. **A**) Representative images of the formation of TIFA oligomers in cells infected with different mutants of *S. flexneri.* HeLa cells expressing TIFA-GFP were left uninfected (mock) or infected for 30 min with indicated strains of *S. flexneri* at MOI 500. pHldE indicates plasmid-encoded HldE. Scale bar, 20 μm**. B**) Infection rate of mutants of the ADP-heptose biosynthetic pathway. HeLa cells were infected at MOI 0.5 with indicated strains and treated with gentamicin. Oligomerization was observed at 4 h post infection with automated microscopy. The fraction of infected cells was analyzed with automated image analysis. Data correspond to the mean +/-SD of 3 independent experiments. **C**) Quantification of TIFA-GFP oligomerization after 30 min of infection with the mutants of the ADP-heptose biosynthetic pathway shown in A. Data correspond to the mean +/-SD of 3 independent experiments. **D**) Quantification of TIFA-GFP oligomerization after 4 h of infection with the *S. flexneri* mutants of the ADP-heptose biosynthetic pathway. Cells were infected for 30 min and treated with gentamicin. Oligomerization was observed at 4 h post infection with automated microscopy. The fraction of infected cells was analyzed with automated image analysis. Data correspond to the mean +/-SD of 3 independent experiments. **E**) Quantification of IL-8 production 4 h post infection with the *S. flexneri* mutants of the ADP-heptose biosynthetic pathway. Cells were infected as in D and treated with monensin. IL-8 was detected by immunofluorescence and quantified by automated image analysis. Data correspond to the mean +/-SD of independent experiments. For comparison between mock and each infected condition (C, D, E) or Δ*hldE* and double-complemented mutant (dash bar), statistical significance was assessed using unpaired, two-tailed Student’s *t*-test, p*<0.05, p**<0.005, p***<0.0005, non-significant (ns).

In conclusion, our data showed that βHBP can only trigger delayed inflammatory signaling, and that it has a marginal contribution in controlling the IL-8 response during *S. flexneri* infection. The delay in the response to βHBP suggests that it has to be intracellularly processed to induce inflammation. βHBP is therefore likely not a true PAMP, which is, by definition, directly sensed via the interaction with a cognate pathogen recognition receptor. Thus, the role of ADP-D/L-β-D-heptose in *S. flexneri* infection challenges our previous results implicating βHBP [3]. The ability of the Δ*gmhB* mutant to induce TIFA oligomerization was the rational basis to conclude that the PAMP inducing TIFA-dependent innate immunity was indeed βHBP. However, this argument is weakened by the possibility that the Δ*gmhB* mutant might only have a partial mutant phenotype. Indeed, the observation that, in contrast to Δ*hldE,* Δ*hldD* and ΔwaaC, the Δ*gmhB* mutant is not hyper-invasive [3], indicated that its LPS was only partially altered and that ADP-D/L-β-D-heptose synthesis was therefore still occurring in these bacteria. A partial phenotype of the Δ*gmhB* mutant was also observed in *E. coli,* and the existence of a yet-unidentified phosphatase activity that compensates for the absence of GmhB was proposed [22]. Given that the Δ*gmhB*-based argument was also used in other studies [11,12], the direct contribution of βHBP has been likely more widely overestimated. The identification of ADP-heptose as a PAMP of *S. flexneri* infection opens a new avenue to understand how inflammation is regulated at the level of the intestinal epithelium in shigellosis. The mechanism of ADP-heptose sensing will have to be thoroughly characterized in space and time during infection. At the time of finalizing this manuscript, Zhou et al. elegantly reported that the kinase ALPK1 is the cytosolic immune receptor for ADP-heptose [23]. More work is required to identify the respective contribution of ADP-D-β-D-heptose and ADP-L-β-D-heptose, to understand how these metabolites are delivered in cells during infection, and in particular, whether they are injected by the type III secretion apparatus of *S. flexneri*. Given that ADP-D/L-β-D-heptose are broadly present in bacteria, our study opens also new perspectives to elucidate the molecular mechanisms regulating innate immunity in infections by other severe human pathogens, including *H. pylori, S. typhimurium and Pseudomonas aeruginosa*. Beyond infections, it will also contribute to characterize new molecular cross-talks between bacteria of the microbiota, the intestinal epithelium and the immune system that regulate intestinal homeostasis.

## MATERIALS AND METHODS

### Cells

HeLa cells (American Type Culture Collection) were cultured in Dulbecco’s modified Eagle’s (DMEM) medium supplemented with 10% FCS and 2 mM Glutamax-1 (complete growth medium). HeLa cells stably expressing TIFA-GFP were obtained after transfection of pEGFP-C1 plasmid encoding TIFA-GFP and geneticin selection at 1 mg.mL^−1^.

### Bacterial strains and generation of mutants

Wild-type (wt), Δ*hldE* and Δ*waaC* M90T *S. flexneri* strains have been previously described [3]. The Δ*hldD* mutant was generated by allelic exchange using a modified protocol of lambda red-mediated gene deletion [24]. Briefly, the kanamycin cassette of the pkD4 plasmid was amplified by PCR with the primers listed in Table 1. The purified PCR product was electroporated into the wt strain expressing the genes for lambda red recombination from the pKM208 plasmid. Recombinants were selected on TSB plates containing 50 μg.mL^−1^ of kanamycin. Single colonies were screened by PCR. *S. flexneri* M90T expressing the *hldA* gene from *N. meningitidis* was previously described [3]. *S. flexneri* M90T expressing the *hldC* gene from *N. meningitidis* was generated as follows. *hldC* was amplified by PCR from a bacterial lysate with the primers listed in Table S1. After gel purification (Macherey-Nagel), the PCR product was digested with EcoRI and HindIII, and ligated into an EcoRI/HindIII-digested pHSG396 plasmid, a derivative of pUC-type of plasmid harboring a chloramphenicol resistance cassette (Clontech). The ligation product was used to transform Top10 *E. coli*. pHSG396-HldC was purified and used to electroporate Δ*hldE and* Δ*hldE* expressing *hldA* strains of S. *flexneri* M90T. All strains were transformed with pCLR7 and constitutively expressed dsRed protein (kind gift from D. Bumann, Biozentrum, Basel, Switzerland).

**Table 1:**
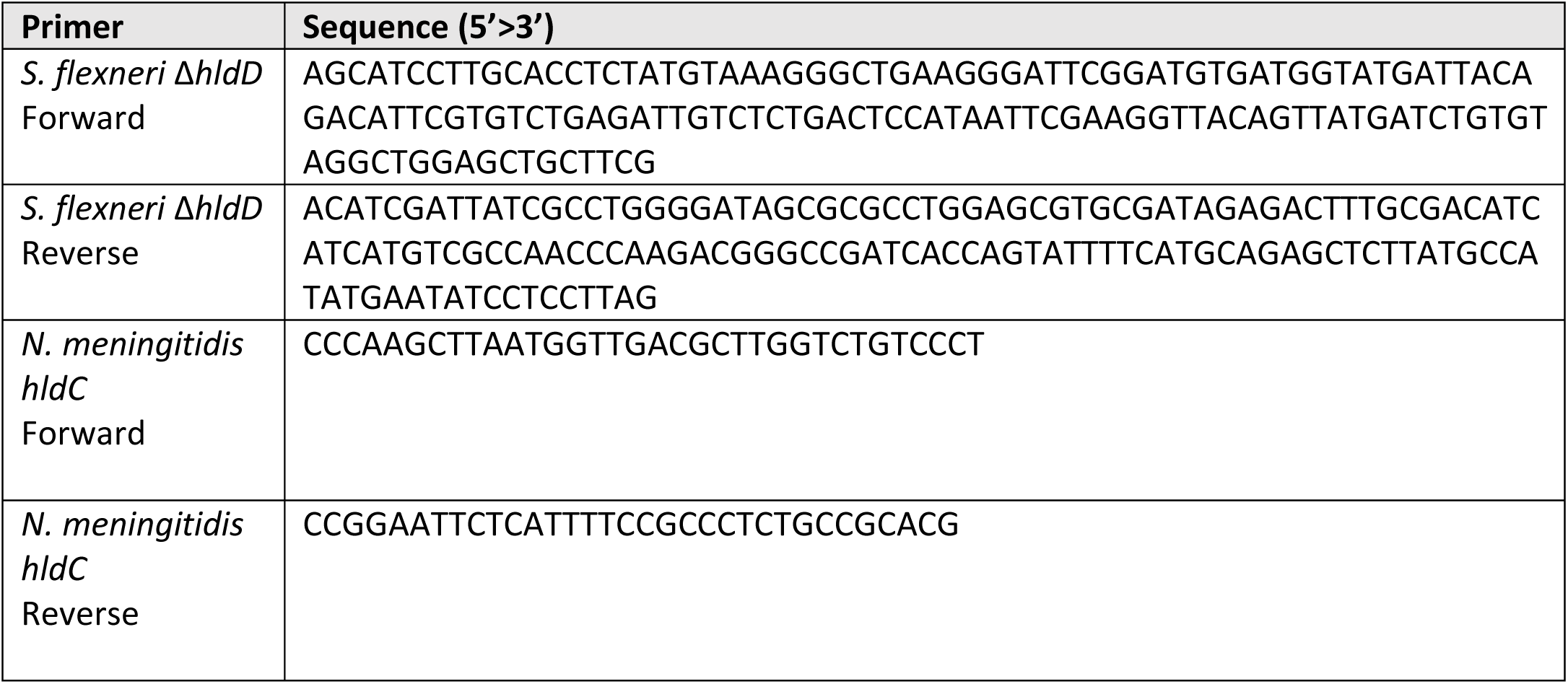
Table listing the primers used in this study.

### Preparation of bacterial lysates

Bacteria from overnight cultures were washed in PBS and resuspended at 10^10^ bacteria.mL^−1^. Once transferred into cryotubes, they underwent through two freeze-thaw cycles in liquid nitrogen. Bacterial debris were removed by centrifugation at 13 000 rpm for 20 min. Lysates were stored at −80 °C.

### Cell treatments

Cells were seeded in 96-well plates the day before the experiment. For treatment, they were washed in permeabilization buffer containing 100 mM KCl, 3 mM MgCl2, 50 mM Hepes, 0.1 mM DTT, 85 mM sucrose, 0.2% BSA and 0.1 mM ATP. They were then incubated with digitonin (5 μg.mL^−1^) alone or a mixture of digitonin and indicated lysates for 30 min in this same buffer. They were then either fixed with 4% PFA or washed to remove digitonin and incubated for 5.5 h in DMEM supplemented with 1% FCS. For IL-8 measurements, cells were treated with 50 μM monensin at 3 hours.

### Infections

For infection experiments with *S. flexneri,* bacteria were used in exponential growth phase. Cells, seeded in 96-well plates, were infected at indicated MOIs in DMEM supplemented with 10 mM Hepes, 2 mM glutamax-1 and 1% FCS. After adding bacteria, plates were centrifuged for 3 min at 1200 rpm and placed at 37 °C for the indicated time periods. Extracellular bacteria were killed by gentamicin (100 μg/mL) and infection was stopped by adding 4% PFA.

### IL-8 measurements

The production of IL-8 was measured by immunofluorescence 6 h after cell treatments or 4 h post infection. In order to trap IL-8 intracellularly, monensin (50 μM) was added after 3 h. After fixation in 4% PFA, cells were stained with an anti-human IL-8 antibody in 0.2% saponin in PBS (BD Pharmingen, San Jose, USA) and an Alexa 647-conjugated secondary antibody (Invitrogen, Carlsbad, USA). IL-8 was quantified by automated image analysis (see below). In parallel, cells were stained with Hoechst (Invitrogen) to visualize nuclei.

### Automated microscopy and image analysis

Images were acquired with an ImageXpress Micro (Molecular devices, Sunnyvale, USA). Each data point correspond to triplicate wells and more than 6 images were taken per well. Image analysis was performed using the custom module editor (CME) of MetaXpress. For IL-8, measurements were performed as previously described [3]. Briefly, cell nuclei were identified by the “Autofind blobs” function. Each nucleus was extended by 6 pixels to define the cell mask in which IL-8 signals were quantified. They were detected with the “Keep marked object” function of CME based on minimal/maximal size requirements and intensity threshold. For the oligomerization of TIFA, foci were detected by applying the Find blobs function based on size and intensity above background parameters. Cell nuclei were identified with the “Find round objects” module. Cells were defined by extending each nucleus by 10 pixels with the “Grow objects without touching” function and TIFA punctates were quantified within this mask. For infection measurements, nuclei and cells were segmented as above. Bacteria were detected by the “Find blobs “function based on size and intensity above background parameters. For IL-8, TIFA oligomerization and infection measurements, the fraction of cells respectively containing IL-8, TIFA oligomers or bacteria was extracted, and used for quantification.

### Chemical synthesis of βHBP

#### General Information

Thin Layer Chromatography (TLC) was performed on precoated slides of Silica Gel 60 F254 (Merck). Detection was effected when applicable, with UV light, and/or by charring with orcinol (35 mM) in 4 N aqueous (aq.) sulfuric acid and ethanol (95:5). Flash chromatography was performed by elution from columns of Silica Gel 60 (particle size 0.040-0.063 mm). Analytical reverse phase high performance liquid chromatography (RP-HPLC) was performed by elution from a C18 X-Bridge Waters Amide column (4.6×250 mm), using CH3CN in 5 mM aq. ammonium formate (0→3%) at 0.5 mL.min^−1^ and detecting at 230 nm. Nuclear Magnetic Resonance (NMR) spectra were recorded at 30 °C for solutions in CDCl3 or D2O (400 MHz for ^1^H, 100 MHz for ^13^C). Residual CHCl3 (7.28 ppm for ^1^H and 77.0 ppm for ^13^C), and HOD (4.79 ppm) were used as internal references for solutions in CDCl3 and D2O, respectively. Proton-signal assignments were made by first-order analysis of the spectra and analysis of 2D ^1^H-^1^H correlation maps (COSY). Of the two magnetically non-equivalent geminal protons at C-7, the one resonating at lower field is denoted H-7a, and the one at higher field is denoted H-7b. Carbon-signal assignments were supported by 2D ^13^C-^1^H correlations maps (HSQC) and analysis of the DEPT and ^13^C spectra. Interchangeable assignments are marked with an asterisk. Electrospray Ionisation-Time Of Flight (ESI-TOF) mass spectra were recorded in the positive-ion mode using a 1:1 CH3CN/water containing 0.1% formic acid ESI-TOF spectrometer-solution. Anhydrous (anhyd.) dichloromethane (DCM), 1,2-dichloroethane (DCE), tetrahydrofuran (THF), *N,N*-dimethylformamide (DMF) and toluene (Tol), sold on molecular sieves (MS) were used as such. Reactions requiring anhyd. conditions were performed under an Argon atmosphere.

#### Synthetic protocols and analytical data

**Phenyl 2,3,4-tri-*O*-benzyl-1-thio-α-D-mannopyranoside (3)**. To a solution of thioglycoside **2** [18] (15.0 g, 55.1 mmol, 1 equiv.) and imidazole (11.3 g, 165.2 mmol, 3 equiv.) in anhyd. THF (110 mL) was added triisopropylsilyl chloride (TIPSCl) (12.4 mL, 58.8 mmol, 1.05 equiv.) dropwise for 15 min at 0 °C. The reaction mixture was stirred at 0 °C for 20 min before warming up to room temperature for 2.5 h, at which time TLC showed the disappearance of the starting material (DCM/MeOH 8:2, Rf = 0.71) and the presence of a less polar compound (Tol/EtOAc 3:7, Rf = 0.6). An additional amount of TIPSCl (0.08 mL, 0.40 mmol, 0.05 equiv.) was added and after 5 h, the reaction was then quenched with aq. saturated (sat.) NH4Cl. The phases were separated, the aq. layer was extracted with EtOAc and the organic phases were washed with brine. The organic phase was dried over Na2SO4, filtered, evaporated under reduced pressure and the crude was used as such in the next step.

To a solution of the crude triol (23.6 g, 55.1 mmol, 1 equiv.) and benzyl bromide (BnBr, 23.6 mL, 198.4 mmol, 3.6 equiv.) in anhyd. DMF (306 mL) was added NaH (60% in mineral oil, 13.2 g, 330.6 mmol, 6 equiv.) portionwise for 15 min at 0 °C. The reaction mixture was allowed to warm up to rt overnight, at which time TLC showed the disappearance of the starting material (Tol/EtOAC 3:7, Rf = 0.6) and the presence of a less polar compound (Cyclohexane/EtOAc 9:1, Rf = 0.75). The reaction was then quenched with MeOH at 0 °C and volatiles were evaporated under reduced pressure. EtOAc and H_2_O were added and the aq. layer was extracted with EtOAc. The organic phases were washed successively with aq. sat. NH4Cl and brine, dried over Na_2_SO_4_, filtered, evaporated under reduced pressure and the crude was used as such in the next step.

To a solution of the silyl ether (38.5 g, 55.1 mmol, 1 equiv.) in anhyd. THF (306 mL), was added tetrabutylammonium fluoride (TBAF, 110 mL, 1 M in THF, 110 mmol, 2 equiv.) at rt. The reaction mixture was stirred for 3 h at which time TLC showed the disappearance of the starting material (Tol/EtOAc 7:3, Rf = 0.9) and the presence of a more polar compound (Tol/EtOAc 7:3, Rf = 0.6). The reaction was quenched with aq. sat. NH4Cl. The phases were separated, the aq. layer was extracted with EtOAc and the organic phases were washed successively with aq. sat. NH4Cl and brine, dried over Na2SO4, filtered and evaporated under reduced pressure. The crude was purified by column chromatography on silica gel using Cyclohexane/EtOAc (100:0 to 80:20) to afford alcohol **3** as a yellow oil (28.0 g, 93% over 3 steps). (ESI^+^)-HRMS: Calcd for C33H38NO5S [M+NH4]^+^ *m/z:* 560.2471, Found: 560.2485. Other analytical data were similar to those published [13].

### Phenyl 2,3,4-tri-*O*-benzyl-6,7-dideoxy-1-thio-α-D-*manno*-hept-6-enopyranoside

**(4)** [19]. To a solution of alcohol **3** (15.0 g, 276.4 mmol, 1 equiv.) in anhyd. DCM (2.0 mL) stirred at 0 °C were added successively DMSO (9.8 mL, 138.2 mmol, 5 equiv.), (DIPEA) (14.5 mL, 82.9 mmol, 3 equiv.) and SO3·Py (13.2 g, 82.9 mmol, 3 equiv.). The reaction mixture was stirred at 0 °C for 15 min and was then allowed to warm up to rt for 4 h. At this time, the reaction was quenched with aq. sat. NH4Cl. The phases were separated, the aq. layer was extracted with DCM and the organic phases were washed successively with aq. sat. NH4Cl and brine, dried over Na2SO4, filtered and evaporated under reduced pressure. The crude was used as such in the next step.

To a suspension of Ph_3_PMeBr (52.3 g, 146.5 mmol, 5.3 equiv.) in anhyd. THF (420 mL) stirred at 0 °C was added *n*-BuLi (33.2 mL, 82.9 mmol, 3 equiv.) dropwise. The yellow solution was warmed up to rt for 1 h. Then, a solution of the obtained aldehyde (14.9 g, 276 mmol, 1 equiv.) in anhyd. THF (10 mL) was added dropwise at −78 °C. The reaction mixture was allowed to warm up to rt overnight, at which time TLC showed the disappearance of the starting material (Tol/EtOAc 8:2, Rf = 0.65) and the presence of a less polar compound (Cyclohexane/EtOAc 8:2, Rf = 0.9). The reaction was quenched with aq. sat. NH_4_Cl. The phases were separated, the aq. layer was extracted with EtOAc and the organic phases were washed successively with aq. sat. NH_4_Cl and brine, dried over Na_2_SO_4_, filtered and evaporated under reduced pressure. The crude was purified by column chromatography on silica gel using Tol/EtOAc (100:0 to 95:5) to afford heptoside **4** as a yellow oil (13.6 g, 85%). (ESI^+^)-HRMS: Calcd for C_34_H_38_NO_4_S [M+NH4]^+^ *m/z:* 556.2521, Found: 556.2527. Other analytical data were similar to those published [13].

**2,3,4,6-Tetra-*O*-benzyl-D-*glycero*-α-D-*manno*-heptopyranose (5).** To a solution of phenyl 2,3,4,6-tetra-*O*-benzyl-1-thio-D-*glycero*-α-D-*manno*-heptopyranoside [13] (1.0 g, 1.51 mmol, 1 equiv.) in a 9:1 mixture of acetone/H2O (15 mL) was added *N*-bromosuccinimide (NBS) (537 mg, 3.02 mmol, 2 equiv.) at 0 °C under Ar and in the dark. The orange mixture was stirred at 0 °C for 15 min then was warmed up to rt. After stirring for 1.5 h, an additional amount of NBS (1 equiv.) was added to the colourless solution at 0 °C. The mixture was stirred overnight at rt, at which time TLC showed the disappearance of the starting material (Tol/EtOAc 8:2, Rf = 0.7) and the presence of a more polar compound (Tol/EtOAc 8:2, Rf = 0.2). The reaction was quenched with aq. sat. Na2S2O3. The phases were separated, the aq. layer was extracted with EtOAc and the organic phases were washed successively with aq. sat. NaHCO3 and brine, dried over Na_2_SO_4_, filtered and evaporated under reduced pressure. The crude was purified by column chromatography on silica gel using Tol/EtOAc (100:0 to 50:50) to afford heptopyranose **5** as a white wax (822 mg, 95%). ^1^H NMR (400 MHz, CDCl_3_) δ 7.38 – 7.17 (m, 20H, H-Ph), 5.18 (brs, 1H, H-1), 4.92 (d, *J* = 11.1 Hz, 1H, H-C<u>H</u>_2_Ph), 4.77 (d, *J* = 12.2 Hz, 1H, H-C<u>H</u>_2_Ph), 4.71 (d, *J* = 12.0 Hz, 1H, H-C<u>H</u>_2_Ph), 4.64 (s, 2H, H-C<u>H</u>_2_Ph), 4.64 (d, *J* = 12.3 Hz, 1H, H-C<u>H</u>_2_Ph), 4.59 (d, *J* = 11.1 Hz, 1H, H-C<u>H</u>_2_Ph), 4.55 (d, *J* = 12.0 Hz, 1H, H-C<u>H</u>2Ph), 4.13 (brd, *J* = 8.6 Hz, 1H, H-5), 4.02 – 3.95 (m, 2H, H-3, H-4), 3.80 (tapp, *J* = 2.2 Hz, 1H, H-2), 3.80 – 3.73 (m, 2H, H-6, H-7a), 3.72 – 3.65 (m, *J* = 14.3, 7.6 Hz, 1H, H-7b), 3.45 (brs, 1H, O<u>H</u>). ^13^C NMR (100 MHz, CDCl_3_) δ 138.3 (Cq-Ph), 138.3 (Cq-Ph), 138.2 (Cq-Ph), 138.1 (Cq-Ph), 128.6 – 127.6 (20C-Ph), 92.5 (C-1), 79.9 (C-3*), 78.1 (C-6), 75.1 (C-2), 74.7 (C-4*), 74.7 (C-<u>C</u>H2Ph), 72.8 (C-<u>C</u>H_2_Ph), 72.7 (C-5), 72.1 (C-<u>C</u>H_2_Ph), 71.8 (C-<u>C</u>H2Ph), 61.8 (C-7). (ESI^+^)-HRMS: Calcd for C35H_42_O_7_N [M+NH4]^+^ *m/z:* 588.2961, Found: 588.2979.

### D-*Glycero*-β-D-*manno*-heptopyranose 1,7-bisphosphate (βHBP)

To a solution of dibenzyl 2,3,4,6-tetra-*O*-benzyl-7-di(benzyloxy)phosphoryl-D-*glycero*-β-D-*manno*-heptopyranosyl phosphate (70 mg, 64 µmol, 1 equiv.) in a 3:1 mixture of 1,4-dioxane/H2O (4 mL) was added Pd(OH)_2_/C (20%, 20 mg) at rt. The reaction mixture was stirred for 16 h under H_2_ (1 atm). The reaction mixture was degassed, filtered on 0.2 µm filter and washed with H2O. The filtrate was adjusted to pH = 7.0 with Na^+^ Dowex 50 and stirred for 30 min. The resin was filtered off and the filtrate was lyophilized to give βHBP (sodium salt form). (ESI^−^)-HRMS: Calcd for C_7_H_15_O_13_P_2_ [M-H]^−^ *m/z:* 368.9988, Found: 368.9974. tr (RP-HPLC): 3.69 min. Other analytical data corresponded those published [13].

## ACKNOWLEGEMENTS

We gratefully acknowledge financial support from the Agence Nationale pour la Recherche (grants n° ANR-14-ACHN-0029-01 and ANR-17-CE15-0006 including postdoctoral fellowships to DGW and JC) and from Institut Pasteur including a postdoctoral fellowship to JC and a master fellowship to LT. We thank Catherine Guerreiro (UCB) for her assistance with HPLC analysis and Frédéric Bonhomme (CNRS UMR3523) for assistance with HRMS measurements and NMR spectroscopy.

## AUTHOR CONTRIBUTIONS

CA, LAM, JC, DGW and ASD designed research. DGW, ASD, JC and LT performed research and analyzed data. HR performed research. CA, LAM and JC wrote the manuscript.

## CONFLICT OF INTEREST

We have no conflict of interest to declare.

## REFERENCES

1. Gaudet RG, Sintsova A, Buckwalter CM, Leung N, Cochrane A, Li J, Cox AD, Moffat J, Gray-Owen SD (2015) INNATE IMMUNITY. Cytosolic detection of the bacterial metabolite HBP activates TIFA-dependent innate immunity. Science 348: 1251–1255.

2. Tettelin H, Saunders NJ, Heidelberg J, Jeffries AC, Nelson KE, Eisen JA, Ketchum KA, Hood DW, Peden JF, Dodson RJ, et al. (2000) Complete genome sequence of Neisseria meningitidis serogroup B strain MC58. Science 287: 1809–1815.

3. Milivojevic M, Dangeard A-S, Kasper CA, Tschon T, Emmenlauer M, Pique C, Schnupf P, Guignot J, Arrieumerlou C (2017) ALPK1 controls TIFA/TRAF6-dependent innate immunity against heptose-1,7-bisphosphate of gram-negative bacteria. PLoS Pathog 13: e1006224.

4. Valvano MA, Marolda CL, Bittner M, Glaskin-Clay M, Simon TL, Klena JD (2000) The rfaE gene from Escherichia coli encodes a bifunctional protein involved in biosynthesis of the lipopolysaccharide core precursor ADP-L-glycero-D-manno-heptose. J Bacteriol 182: 488–497.

5. Taylor PL, Sugiman-Marangos S, Zhang K, Valvano MA, Wright GD, Junop MS (2010) Structural and kinetic characterization of the LPS biosynthetic enzyme D-alpha, beta-D-heptose-1,7-bisphosphate phosphatase (GmhB) from Escherichia coli. Biochemistry 49: 1033–1041.

6. Takatsuna H, Kato H, Gohda J, Akiyama T, Moriya A, Okamoto Y, Yamagata Y, Otsuka M, Umezawa K, Semba K, et al. (2003) Identification of TIFA as an adapter protein that links tumor necrosis factor receptor-associated factor 6 (TRAF6) to interleukin-1 (IL-1) receptor-associated kinase-1 (IRAK-1) in IL-1 receptor signaling. J Biol Chem 278: 12144–12150.

7. Li J, Lee GI, Van Doren SR, Walker JC (2000) The FHA domain mediates phosphoprotein interactions. J Cell Sci 113 Pt 23: 4143–4149.

8. Huang C-CF, Weng J-H, Wei T-YW, Wu P-YG, Hsu P-H, Chen Y-H, Wang S-C, Qin D, Hung C-C, Chen S-T, et al. (2012) Intermolecular binding between TIFA-FHA and TIFA-pT mediates tumor necrosis factor alpha stimulation and NF-?B activation. Mol Cell Biol 32: 2664–2673.

9. Ea C-K, Sun L, Inoue J-I, Chen ZJ (2004) TIFA activates IkappaB kinase (IKK) by promoting oligomerization and ubiquitination of TRAF6. Proc Natl Acad Sci U S A 101: 15318–15323.

10. Zimmermann S, Pfannkuch L, Al-Zeer MA, Bartfeld S, Koch M, Liu J, Rechner C, Soerensen M, Sokolova O, Zamyatina A, et al. (2017) ALPK1-and TIFA-Dependent Innate Immune Response Triggered by the Helicobacter pylori Type IV Secretion System. Cell Rep 20: 2384–2395.

11. Gall A, Gaudet RG, Gray-Owen SD, Salama NR (2017) TIFA Signaling in Gastric Epithelial Cells Initiates the cag Type 4 Secretion System-Dependent Innate Immune Response to Helicobacter pylori Infection. mBio 8:.

12. Stein SC, Faber E, Bats SH, Murillo T, Speidel Y, Coombs N, Josenhans C (2017) Helicobacter pylori modulates host cell responses by CagT4SS-dependent translocation of an intermediate metabolite of LPS inner core heptose biosynthesis. PLoS Pathog 13: e1006514.

13. Liang L, Vincent SP (2017) Synthesis of d-glycero-d-manno-heptose 1,7-bisphosphate (HBP) featuring a β-stereoselective bis-phosphorylation. Tetrahedron Lett 58: 3631–3633.

14. Sauvageau J, Bhasin M, Guo CX, Adekoya IA, Gray-Owen SD, Oscarson S, Guazzelli L, Cox A (2017) Alternate synthesis to d-glycero-ß-d-manno-heptose 1,7-biphosphate. Carbohydr Res 450: 38–43.

15. Inuki S, Aiba T, Kawakami S, Akiyama T, Inoue J-I, Fujimoto Y (2017) Chemical Synthesis of d-glycero-d-manno-Heptose 1,7-Bisphosphate and Evaluation of Its Ability to Modulate NF-?B Activation. Org Lett 19: 3079–3082.

16. Borio A, Hofinger A, Kosma P, Zamyatina A (2017) Chemical synthesis of the innate immune modulator – bacterial d-glycero-ß-d-manno-heptose-1,7-bisphosphate (HBP). Tetrahedron Lett 58: 2826–2829.

17. Patent WO2012073214A2 -Recherche Google.

18. Crich D, Chandrasekera NS (2004) Mechanism of 4,6-O-benzylidene-directed beta-mannosylation as determined by alpha-deuterium kinetic isotope effects. Angew Chem Int Ed Engl 43: 5386–5389.

19. Brimacombe JS, Kabir AKM (1986) Convenient syntheses of l-glycero-d-manno-heptose and d-glycero-d-manno-heptose. Carbohydr Res 152: 329–334.

20. Bennett BD, Kimball EH, Gao M, Osterhout R, Van Dien SJ, Rabinowitz JD (2009) Absolute metabolite concentrations and implied enzyme active site occupancy in Escherichia coli. Nat Chem Biol 5: 593–599.

21. Kneidinger B, Marolda C, Graninger M, Zamyatina A, McArthur F, Kosma P, Valvano MA, Messner P (2002) Biosynthesis Pathway of ADP-l-glycero-ß-d-manno-Heptose in Escherichia coli. J Bacteriol 184: 363–369.

22. Nakao R, Ramstedt M, Wai SN, Uhlin BE (2012) Enhanced Biofilm Formation by Escherichia coli LPS Mutants Defective in Hep Biosynthesis. PLoS ONE 7:.

23. Zhou P, She Y, Dong N, Li P, He H, Borio A, Wu Q, Lu S, Ding X, Cao Y, et al. (2018) Alpha-kinase 1 is a cytosolic innate immune receptor for bacterial ADP-heptose. Nature.

24. Datsenko KA, Wanner BL (2000) One-step inactivation of chromosomal genes in Escherichia coli K-12 using PCR products. Proc Natl Acad Sci U S A 97: 6640–6645.

